# Plastic biodegradation: do *Galleria mellonella* larvae - bio-assimilate polyethylene? A spectral histology approach using isotopic labelling and infrared microspectroscopy

**DOI:** 10.1101/2021.10.08.463624

**Authors:** Agnès Réjasse, Jehan Waeytens, Ariane Deniset-Besseau, Nicolas Crapart, Christina Nielsen-Leroux, Christophe Sandt

## Abstract

Environmental pollution by non-biodegradable polyethylene (PE) plastics is of major concern, thus, organisms capable of bio-degrading PE are required. The larvae of the Greater Wax Moth, *Galleria mellonella* (Gm), were identified as a potential candidate to digest PE. In this study, we tested whether PE was metabolized by Gm larvae and could found in their tissues. We examined the implication of the larval gut microbiota by using conventional and axenic reared insects. First, our study showed that neither beeswax nor PE alone favour the growth of young larvae. We then used Fourier-Transform Infrared Microspectroscopy (µFTIR) to detect deuterium in larvae fed with isotopically labelled food. Perdeuterated molecules were found in most tissues of larvae fed with deuterium labelled oil for 72 hours proving that µFTIR can detect metabolization of 1-2 mg of deuterated food. No bio-assimilation was detected in the tissues of larvae fed with 1-5 mg of perdeuterated PED4 for 72 hours and 19-21 days, but micron sized PE particles were found in the larval digestive tract cavities. We evidenced weak bio-degradation of PE films in contact for 24 hours with the dissected gut of conventional larvae; and in the PED4 particles from excreted larval frass. Our study confirms that Gm larvae can bio-degrade PE but can not necessarily metabolize it.

## INTRODUCTION

Due to inefficient waste collection and long-life, plastics are now a major cause of environmental pollution in land and maritime environments^1^. Development of new biodegradable plastics and new plastic degradation processes are pursued to remediate these problems. Biodegradation offers advantages over other methods (incineration, chemical degradation) as it can be more environmentally friendly, produces less waste, and reduces cost of waste management. Polyethylene (PE), one of the most produced plastics, is considered almost non-biodegradable. Furthermore, to increase its lifetime, PE is generally synthetized with antioxidants and UV-stabilizers^2^. Modest biodegradation rates were reported in the literature by microorganisms from natural microbial communities ^3–6^.

Another potential plastic biodegradation method reported in the literature is the use of insect larvae or their commensal gut microorganisms^7–13^. Several recent studies reported degradation of PE by the caterpillars of the Greater Wax Moth *Galleria mellonella* (Gm)^11,12,14,15,16^. There is a rational motivation for the use of these larvae. The metabolic pathways involved in the degradation of long-chain hydrocarbons (like long-chain fatty acids) are expected to play an important role in the degradation of PE that is composed of a long aliphatic chain. Since,, Gm larva feeds on and metabolizes long-chain hydrocarbons from beeswax^17^, it may also potentially metabolize PE. If this is the case, the role of gut enzymes and the gut microbiota should be assessed. However, involvement of the larval gut microbiota may be questionable. Recently, Kong et al.^14^ reported PE biodegradation independent of the intestinal microbiota while Ren^12^, Cassone^16^ and Lou^18^ described implication of various species from the Gm gut microbiota. In addtion the original study^11^ reporting the degradation of PE by the Gm larva was criticized on several methodological pointsby Weber et al.^19^,. Indeed, the approach used to investigate the potential of the larvae to metabolize PE in those studies presents several issues: gut residues on the PE films are often misinterpreted for PE oxidation; the ingestion of PE does not imply metabolization of the polymer, PE metabolization by the larvae was never demonstrated. This and the controversial results about microbiota involvement make it necessary to further investigate whether Gm larvae and/or their microbiota can really biodegrade and metabolize PE.

In this study, we present a methodology capable of detecting the eventual metabolization of PE by Gm larvae. First, we tested whether PE can be used as an energy source and if it provided nutritional value for the Gm larvae. Then, we used Fourier-Transform Infrared microspectroscopy (µFTIR) to perform hyperspectral imaging of cryo-sections of the whole larvae and we evaluated the capability of Gm larvae to bio-assimilate PE as well as its integration in the larval tissues. We developed an original protocol using polyethylene isotopically labelled with deuterium (PED4) to detect if deuterated molecules were metabolised in the Gm larval tissue after perdeuterated PE ingestion. Indeed, based on its infrared spectrum, the ––CH_2_–– peak from PE cannot be distinguished from the ––CH_2_–– peak from lipids in larva tissues at low concentrations whereas ––CD_2_–– found in deuterated PE has specific absorption peaks in the infrared transparency window of the tissues and could be easily identified in the larvae. The large shift in peak positions between C-H and C-D is caused by the larger atomic mass of deuteriumcausing a strong shift in the vibration frequency of the C–D bonds. This method is one of the most relevant to reveal metabolization. Furthermore, PED4 and regular PE exhibit similar chemical and physical properties and their biodegradation products are expected to be identical. After metabolization, PED4 should be found as deuterated molecules containing ––CD_2_–– moieties, presumably in tissues containing long aliphatic chains molecules. Detecting ––CD_2_–– groups in the tissues of the larvae should be a direct indication of PE metabolization. The sensitivity of the method was evaluated by feeding experiments with deuterated oil. The implication of the gut microbiota in the biodegradation process was evaluated using both conventional and axenically reared larvae (without microbiota). The PE nutritional value was evaluated by following larval growth with different diets at two development stages.

## MATERIALS AND METHODS

### PE materials

Low density PE supermarket bags were used for insect feeding and growth experiments. High density PE bags were used for assessing the oxidation of PE in contact with the dissected Gm larva gut. Perdeuterated PE powder (PED4) was purchased from Medical Isotopes Inc (Pelham, NH, USA). The PEs used in this study were characterized by High-Temperature Gel Permeation Chromatography (HT-GPC) by the Peakexpert company (Tours, France). HT-GPC was performed at 150°C in stabilized trichlorobenzene on Agilent Mixed-B columns. Columns were calibrated with polystyrene references. The molecular weights were found as follows:

| Sample | Mn<br>(g/mol) | Mw<br>(g/mol) | Mz<br>(g/mol) |
| --- | --- | --- | --- |
| LDPE bags | 40,737 | 249,031 | 679,051 |
| HDPE bags | 34,151 | 641,115 | 4,277,300 |
| polyethylene-D4 | 139,715 | 821,776 | 3,354,440 |

### Insect rearing and feeding

*Gm* larvae were produced on site in the insectarium at INRAE Micalis Institute at Jouy-en-Josas. Gm eggs were hatched at 27 °C, and the larvae were reared on beeswax and pollen (La Ruche Roannaise, Roanne, France).

The moths laid eggs on paper that were directly placed on pollen and covered with beeswax in closed aerated plastic boxes. For the axenic larvae, eggs were first sterilized by 10 min exposure on each side with UV light at 254 nm, and fed with gamma-ray sterilized pollen and beeswax in an autoclaved glass jar with aerated lid covered by sterile gauze and carded cotton. The boxes and the jars were placed in a incubator at 27 °C simulating the day-night cycle. In order to verify that the larvae were axenic, 2 larvae were crushed and homogenized with a sterilized pestle in 500 µl sterile physiological water, 100 μL of the suspension were spread on a BHI Petri dish. No bacterial growth was observed after 7 days at 37 °C and no bacterial 16S DNA was found following V3/V4 PCR (For conventional larvae 10^6^ bacteria/larva are found)

### Effect of diet on insect growth

L2-L3 early stage larvae (20 mg each) were placed individually in 12-well plate wells and were either starved or fed *ad libitum* with one of five different diets: beeswax alone, pollen alone, LDPE alone, beeswax + pollen, beeswax + LDPE. Six larvae were used for each diet for a total of 36 larvae. The food uptake was evaluated by weighing the remaining food. The larvae were kept at 27 °C and weighted individually every 2-3 days. The test was continued for 16 days until the larvae fed a beeswax + pollen diet reached stage L6.

A second test was carried out with another batch of L6, (last larval stage /160 mg), 36 larvae (6 for each diet)). The larvae were placed individually in 6-well plates at 27 °C and fed as previously described. The larvae reached the chrysalis stage in 8-10 days; the test was carried-out for an additional 12 days until the adult stage.

The average growth of the 6 larvae and the standard deviations were calculated with the “Origin Pro 2016” software (Origin lab, Northampton, MA).

### Perdeuterated oil feeding

Five conventional and 5 axenic L6 stage larvae were starved for 24 hours before free-feeding for 24 or 72h with pollen soaked with perdeuterated oil (N-hexadecane (C_16_D_34_, 98%) Cambridge Isotope Laboratories, Inc. USA). Larvae fed for 24h ingested 6.7 mg of pollen and 1.6µL of oil (poids ?) and larvae fed for 72h ingested 20 mg of pollen and 4.8 µL of oil.

### Perdeuterated polyethylene (PED4) feeding

PED4 films were prepared either by pressing PED4 powder at 140 °C for 15-30 minutes in an in-house designed pressgiving 30 to 40 µm thick films, or by pressing PED4 powder at room temperature for 1 minute at 15 ton/cm^2^ in a Specac manual hydraulic press (Eurolabo, Paris, France).

Two batches of L6 stage larvae (5 axenic and 5 conventional) were starved for 24h at 27 °C in individual boxes. The larvae were then allowed to feed freely on PED4 films for 3 days with an allowance of at least 0.85 mg of PED4 per day. The larvae were then killed by quick freezing in SnapFrost equipment and cryo-sectioned. The amount of PED4 ingested was evaluated by weighting the PED4 film leftover and feces were collected and stored at −80 °C for further evaluation. Only larvae fed with more than 1 mg of PED4 were analyzed by µFTIR. Three axenic and three conventional larvae were fed for up to 19-21 days with PED4 alternating with pollen diets to allow survival.

### Cryo-sectioning and preparation for hyperspectral IR imaging

The cryo-sections were prepared on the Abridge platform (INRAE, Jouy-en-Josas, France). The larvae were frozen in a SnapFrost system (Excilone, Elancourt, France) in isopentane at −80 °C then stored at −80 °C. Ten and twenty micron sagittal sections were cryo-sectioned at −20 °C with a Shandon FSE cryostat (ThermoFisher Scientific, Courtaboeuf, France). The 20 µm thick sections improved the sensitivity of the detection for the weak C-D peaks but tissue lipid and protein peaks were saturated. - Consecutive sections were made for both kinds of sections and deposited on different slide supports: on StarFrost (Knittelglass, Germany) for immediate cresyl-violet staining, on IR-grade polished IR-transparent CaF_2_ slides (Crystran, Poole, UK) for IR hyperspectral imaging. The sections on CaF_2_ were stored in a desiccator under a continuous flow of nitrogen until analysis.

### Hyperspectral FTIR Imaging

Fourier-Transform Infrared hyperspectral images were recorded at the SMIS beamline on a Cary 620 infrared microscope (Agilent, Courtaboeuf, France) equipped with a 128 x128 pixels Lancer Focal Plane Array (FPA) detector and coupled to a Cary 670 spectrometer. Hyperspectral images were measured in transmission in the standard magnification mode with a 4X/0.2 NA Schwarzschild objective and matching condenser giving a field of view of 2640 × 2640 µm^2^, and a projected pixel size of 20.4 × 20.4 µm^2^. The actual spatial resolution of the images was evaluated by the step-edge method to be approximately 40 µm at 1545 cm^-1^ and 30 µm at 2915 cm^-1^. Mosaics composed of several FPA tiles were recorded to image whole sections.

Hyperspectral images were recorded between 900 and 3900 cm^-1^ at 8 cm^-1^ resolution, with 256 and 128 co-added scans for background and sample respectively.

### Synchrotron Radiation FTIR Microspectroscopy (SR-µFTIR)

SR-µFTIR was performed at the SOLEIL synchrotron facility on the SMIS beamline ^20^. The synchrotron was operated at 500 mA in top-up mode for injections. Spectra and maps of the sections were recorded using Continuum microscopes coupled to Nicolet 8700 or 5700 spectrometers (Thermo Fisher Scientific, Courtaboeuf, France). The microscopes were equipped with 32X/0.65NA Schwarzschild objectives and matching condensers, and liquid-cooled narrow-band MCT/A detectors. Spectra were recorded in transmission mode at 6 cm^-1^ resolution with 16 to 32 scans between 650 and 4000 cm^-1^. The confocal aperture of each microscope was set at 12 × 12 µm^2^.

### Data analysis

Spectral images were computed in ResolutionPro (Agilent) and in Quasar^21,22^. Spectral images were created using the baseline-corrected, integrated areas of the peaks of interest. Lipid and protein distributions were imaged by the C-H stretching peaks of CH_2_ and CH_3_ between 2800 and 3000 cm^-1^ and the amide I band between 1590 and 1705 cm^-1^ respectively. The deuterated PE was detected and imaged by symmetric and asymmetric CD_2_ stretching peaks at 2085 cm^-1^ (2030-2130 cm^-1^) and 2190 cm^-1^ (2165-2230 cm^-1^). The deuterium/protein peak area ratio was computed with protein band area integrated between 1480 and 1720 cm^-1^. K-means clustering of the hyperspectral images and computation of Pearson correlation coefficients between peak-area ratios were performed in Quasar. K-means clustering is a multivariate pattern-recognition method that allows clustering spectra based on their similarity, 10 runs and 300 iterations were used. Water vapour subtraction was performed in Matlab 2016 (MathWorks, Natick, MA) with an in-house script.

## RESULTS

### Nutritional value of PE

To estimate the relative nutritional value of PE as energy source for Gm, we compared the food uptake, weight gain and larval survival for different diets: control (nothing), pollen alone, beeswax alone, beeswax + pollen, LDPE alone, LDPE + pollen. We set up growth experiments with conventional larvae at two different stages (Figure 1A, 1E): young larvae (stage L2-L3, 20 mg per larva) and last instar (stage L6, 160 mg).

**Figure 1.**
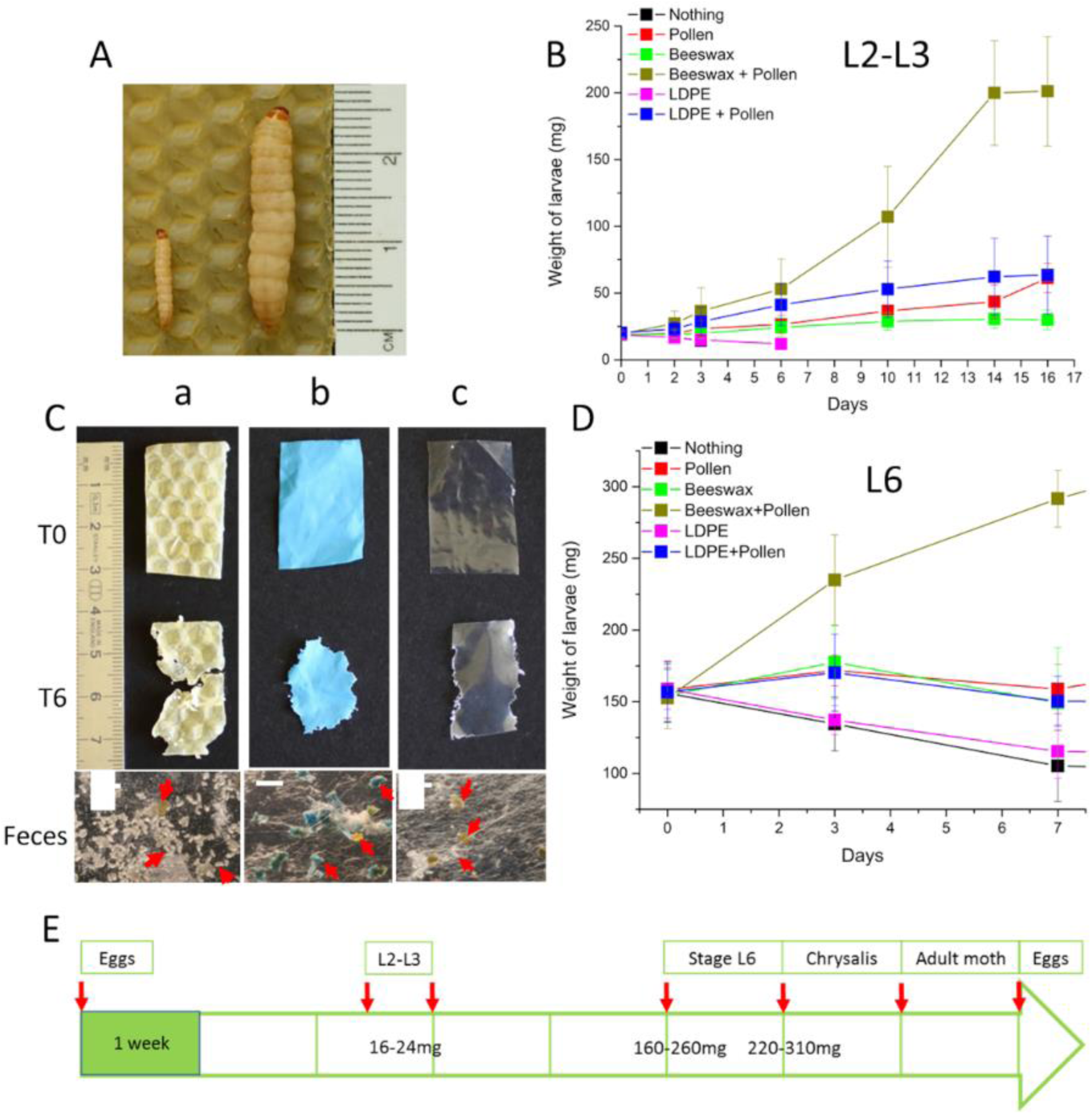
Evaluation of the nutritional value of different diets in Gm. A) Gm larvae on beeswax: L2-L3 stage on the left, L6 stage on the right. B) Growth curve of L2-L3 Gm larvae fed different diets. C) Pictures evidencing the consumption of beeswax or LDPE and the excretion of LDPE in feces. Arrows show the feces among the silk fibers. Scale bars: 2 mm. D) Growth curve of last stage L6 Gm fed different diets. E) Typical Gm larva evolution time scale (weeks) and larval weight (L2 –L3 and L6) when fed with beeswax and pollen. Larvae were reared at 27 °C.

For L2-L3 larvae (Figure 1B) the nibbling of PE was difficult to observe, LDPE consumption was estimated by weighting the remaining PE (Supplementary Table 1). The larval weight uptake was followed up to 16 days. The weight of larvae fed with a pollen-beeswax diet increased 10 times while with a pollen-only diet, it increased 3 times, and 1.3 times with a beeswax-only diet. The pollen-beeswax diet was the optimal condition for Gm larval growth. The control larvae, without food, died after 3 days. Larvae fed with only LDPE lost weight (1.3 fold), did not change growth stage, and exhibited 50% mortality at day 3 and 100% mortality at day 6, probably due to starvation. Larvae fed with both LDPE and pollen did not gain weight compared to larvae fed only with pollen. This indicated that, although they consumed LDPE (Figure 1B), it did not provide energy for growth or survival at early development stages (see Supplementary Table 1 for detailed diet consumption).

For L6 larvae, LDPE consumption was observed directly on PE films as well as residues of colored PE in the excreted feces (Figure 1C),. We followed the growth of L6 stage larvae fed with different diets for 7 days (Figure 1D, and Sup dat Table 2). The results demonstrate that as for the young larvae, the best diet was a combination of beeswax and pollen, since the larvae almost doubled weight in 7 days. Larvae fed with pollen-only diet, wax-only diet or LDPE-pollen diet survived but did not gain weight. Larvae fed with LDPE-only diet lost 25% of their weight as did the control larvae (no food) but all survived and were able to complete metamorphosis into a moth, like in all the other conditions. In average, larvae fed with a pollen-LDPE diet had each consumed 0.48 mg LDPE /day/, larvae fed with LDPE-only diet consumed 0.37 mg LDPE/day/; these rates are similar to those in Lou^18^ (0.60 mg PE/day/larva) but inferior to those reported by Bombelli^11^ (1.84 mg PE/day/larva).

These experiments showed that conventional Gm larvae have eaten PE but no apparent nutritional value was observed nor measured.

### Hyperspectral infrared imaging of larva cryo-sections

Although the larvae did gain weight by eating PE in the aforementioned experiment, it does not prove that Gm larvae or their microbiota could not metabolize small quantities of PE. Therefore, we developed a method based on µFTIR hyperspectral imaging for measuring the chemical tissue composition of the Gm larvae. The ultimate goal was to detect the presence of metabolized PE in the tissue following ingestion with deuterated PE.

First, we set up µFTIR hyperspectral imaging experiment to measure the spectral tissue composition (sugars, proteins and lipids) and if deuterium can be detected in Gm larvae thin-sections in larvae fed with an optimal pollen and wax diet. Hyperspectral infrared images of 10 µm thick cryo-sections of control larvae were recorded (Figure 2A-E). The spectra and peak area used to generate the spectral maps are shown in Figure 2L and Supplementary Figure S1. Representative spectra from different tissues were obtained by classifying all the larval tissue spectra in 4 groups by k-means clustering (Supplementary Figure S1A). All IR absorption spectra from the larval tissues were typical to other biological tissues. They were dominated by the absorption bands of the stretching vibration of O-H and N-H bonds present in sugars and proteins at 3400 and 3300 cm^-1^; C-H peaks from lipids and proteins between 3020 and 2800 cm^-1^, C=O peaks from esterified lipids at 1740 cm^-1^ and carboxylic acids at 1710 cm^-1^; CONH peaks from proteins at 1654 and 1545 cm^-1^, C-H and COOH peaks between 1480 and 1350 cm^-1^; P=O peaks at 1240 and 1080 cm^-1^, C-OH, C-OP and COC and COH peaks from carbohydrates and lipids at 1160, 1150, 1100, 1035 and 1025 cm^-1^. While proteins peaks are generally the most intense peaks in the spectra of most animal tissues, peaks from phospholipids and esterified lipids (C-H, C=O, C-OC, and C-OP) strongly dominated the spectra of larva tissues showing their extremely high lipid concentration. The C=C-H olefinic peak from unsaturated lipids at 3008 cm^-1^ was detected in the lipid rich tissues evidencing a strong lipid unsaturation level. In Figure 2B and 2C, we present respectively the protein and lipid hyperspectral images from the control larva section shown in Figure 2A. Silk glands and some epithelial regions (evidenced by the amide I band at 1650 cm^-1^) appeared like red hotspot in the protein image while fatty tissues (evidenced by esterified lipids by the ester C=O peak at 1740 cm^-1^) appeared red in the lipid image.

**Figure 2.**
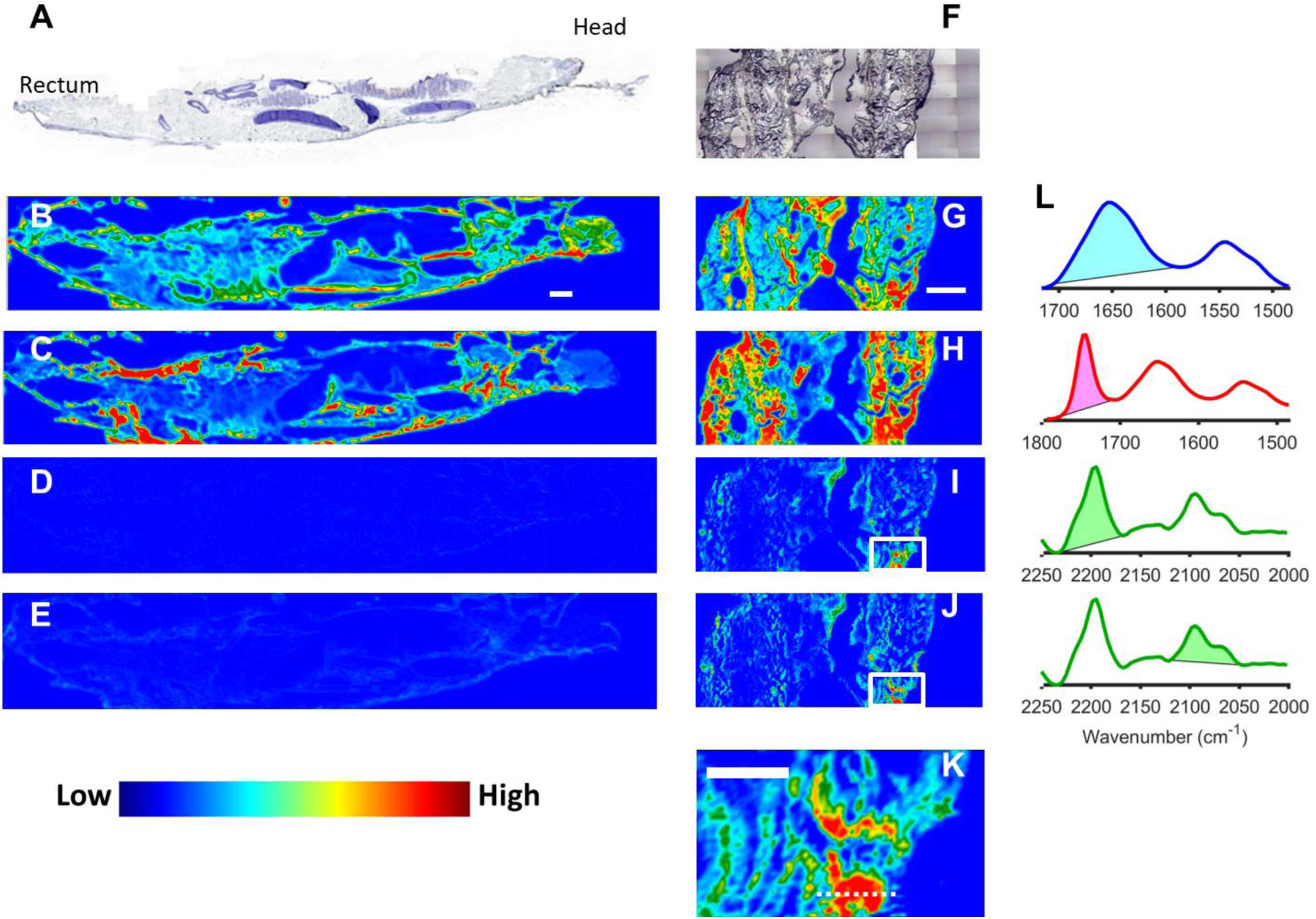
Spectral histology of Gm larvae and detection of C16D34 signal in larva sections. A-E) Micrographs of section of a control larva (4X magnification). A) Bright field of cresyl violet stained section. B-E) Infrared spectral histology maps showing: the distribution of B) proteins, and C) lipids; the absence of C-D peak at D) 2197 cm^-1^, and at E) 2098 cm^-1^. This shows that deuterium is not found in the control larva. F-K) Micrographs of unstained larva fed 72h with C16D34 oil. F) Bright field of the unstained section. G-K) Infrared maps of G) protein distribution, H) lipid distribution. The detection of C-D peaks in the deuterated oil fed larvae is evidenced in I) by the 2197 cm^-1^ C-D peak area from CD2 asymmetric stretching, and in J) by the 2098 cm^-1^ C-D peak area from CD2 symmetric stretching). K) Zoom on a C-D rich region delimited by a box in images I and J. Scale bars: 0. 5 mm. L) The peak area used for plotting the spectral maps, in descending order: protein amide I, lipid ester C=O, symmetrical and asymmetrical C-D stretching peaks of the C_16_D_34_ oil.

Detection of the natural abundance of deuterium in control larva was done by looking at the 2000-2200 cm^-1^ range that contains the strong C-D stretching peaks. The typical C-D peaks are shown in Supplementary Figure S1B in the spectrum of deuterated PED4 (red) and of deuterated oil (green) dominated by the C-D stretching peak at 2197 and 2098 cm^-1^, shown along with the spectrum of a normal PE film (blue) dominated by the methylene (–CH_2_–) peaks at 2914 and 2848 cm^-1^. Since the larval tissue was so rich in lipids, the highest concentration of deuterium might be found in the fatty tissues. Figure 2D and 2E show hyperspectral maps of the larva cryo-section at 2197 and 2098 cm^-1^ respectively, evidencing the absence of detectable C-D peaks in the control larvae. The faint tissue contours observed in Figure 2E arise from baseline drifts caused by IR radiation scattering at the edge of the tissue and not from the C-D peak

### Detection of deuterium in cryo-sections of larva fed with deuterated oil

In order to prove that deuterium could be detected (as C-D bonds) in larvae that had ingested deuterated food, conventional and axenic larvae were fed during 24 and 72 h with C16D34 perdeuterated oil. The oil was mixed with pollen to facilitate its ingestion since larvae would not ingest pure oil. In average each larva had ingested 1.2 mg of oil in 24 h and 3.7 mg in 72 h. The larvae were then cryosectioned and the thin sections were investigated by µFTIR (Figure 2F-K). Weak C-D signal could be detected after 24 h at discrete locations (not shown) but C-D signal was consistently detected in most tissues after 72 h of feeding (Figure 2I, 2J). This suggests that the threshold for consistent detection was around 2-2.5 mg of ingested deuterated food. The C-D signal was detected at discrete locations throughout the sections and appeared stronger in some lipid rich tissues The related spectra can be seen in Supplementary Figure S1C showing the pure C_16_D_34_ oil spectrum and the spectra from the different larval tissues. We found a moderately positive correlation between the distribution of lipids and deuterated molecules with a Pearson correlation coefficient of 0.42 between the C-D and lipid signals. This confirmed that deuterated food assimilation could be detected more sensitively in the fat tissues. Figure 2K shows a zoom on a C-D rich region, the spectra are shown in Supp. Figure S1D. The C-D signal measured with the µFTIR imaging system was low with a maximum of 0.025 A.U. at 2197 cm-1 in the hot spot shown in Figure 2K. We used a confocal microscope coupled to a synchrotron source to improve measurement sensitivity and accuracy (Figure S1E) and measured one order of magnitude higher C-D signal (up to 0.20 A.U.) than with the imaging system. The synchrotron data were then used to examine if the oil was changed upon metabolization: the CD_2_ peak position (sensitive to molecule environment) and the CD_2_/CD_3_ ratio (related to the aliphatic chain length) were determined and compared in the pure oil and in the deuterated fatty tissues. The CD_2_/CD_3_ ratio measured at 24 different positions in the larval tissue varied between 0.72 and 3.6 (mean 2.16) and was different from its value in the pure oil (1.92) showing that the oil was fragmented and CD_2_ moieties were integrated in short and in longer chain lipids. This supported the idea of metabolization of the oil in the larva. The CD_2_ peak positions in pure oil and in the tissues were not significantly different at 2197.1 ± 1.2 cm-1 and 2197.3 ± 0.4 cm-1 respectively, according to a Student T-test. The positions of C-D peaks are sensitive to the conformation of lipids^23^, to their local environment, and to the long range order such as the organization in amorphous gel or liquid crystal. This showed that the deuterated lipids were in similar environments in the pure oil and in the tissues

These results clearly demonstrate that the larvae were able to metabolize and integrate deuterated food into their fatty tissue and that µFTIR hyperspectral imaging is sensitive enough to detect the C-D signal.

### Investigations into cryo-sections of larva fed with deuterated PE

Therefore, next we investigated whether Gm larvae fed with deuterated PE were able to assimilate PE by seeking the appearance of C-D peaks in the larval tissues. Similarly, we recoreded hyperspectral infrared images from sections o from 5 conventional and 5 axenic L6 larvae fed for 3 days with deuterated PE (PED4), and 2 conventional and 2 axenic control larvae fed with non-deuterated PE. For each larva, two cryo-sections, one 10 µm thick and one 20 µm thick, were analysed.. Three additional L3 larvae were fed for 19 days with PED4 alternating with a pollen diet to keep them alive. Each larva ingested in average 2.1 mg of PED4 This corresponds to 0.7 mg PED4/day/larva for L6 larvae. This rate is similar to that of normal LDPE and for those reported in the literature in Lou et al.^18^

The results are presented in Figure 3 which shows visible and IR hyperspectral images of one representative larval cryo-section for each condition. IR hyperspectral images of protein peak at 1650 cm^-1^ and CD_2_ peak at 2197 cm^-1^ are shown. In the control axenic and conventional larvae, no absorption of C-D could be detected in the tissues, as expected, and no C-D peak was detected in the tissues of the 5 conventional and 5 axenic larvae fed for 3 days with PED4. To ensure that it was not due to a sensitivity issue, 20 µm thick sections were studied, giving the same results. In order to further boost the sensitivity of the method another set of conventional and axenic L3 stage larvae was fed during 12 to 19 days with PED4. To make sure that the larvae could survive over 6 days with this low nutritional value diet, the 12- and 19-days periods were cut in periods of 3 days alternating between PED4-only diet and PED4 plus pollen diet. TheseL3 larvae had consumed 3.6 ± 1.1 mg PED4 per larva at the end of the experiment at rates of 0.19 to 0.28 mg PED4/day/larva.

**Figure 3.**
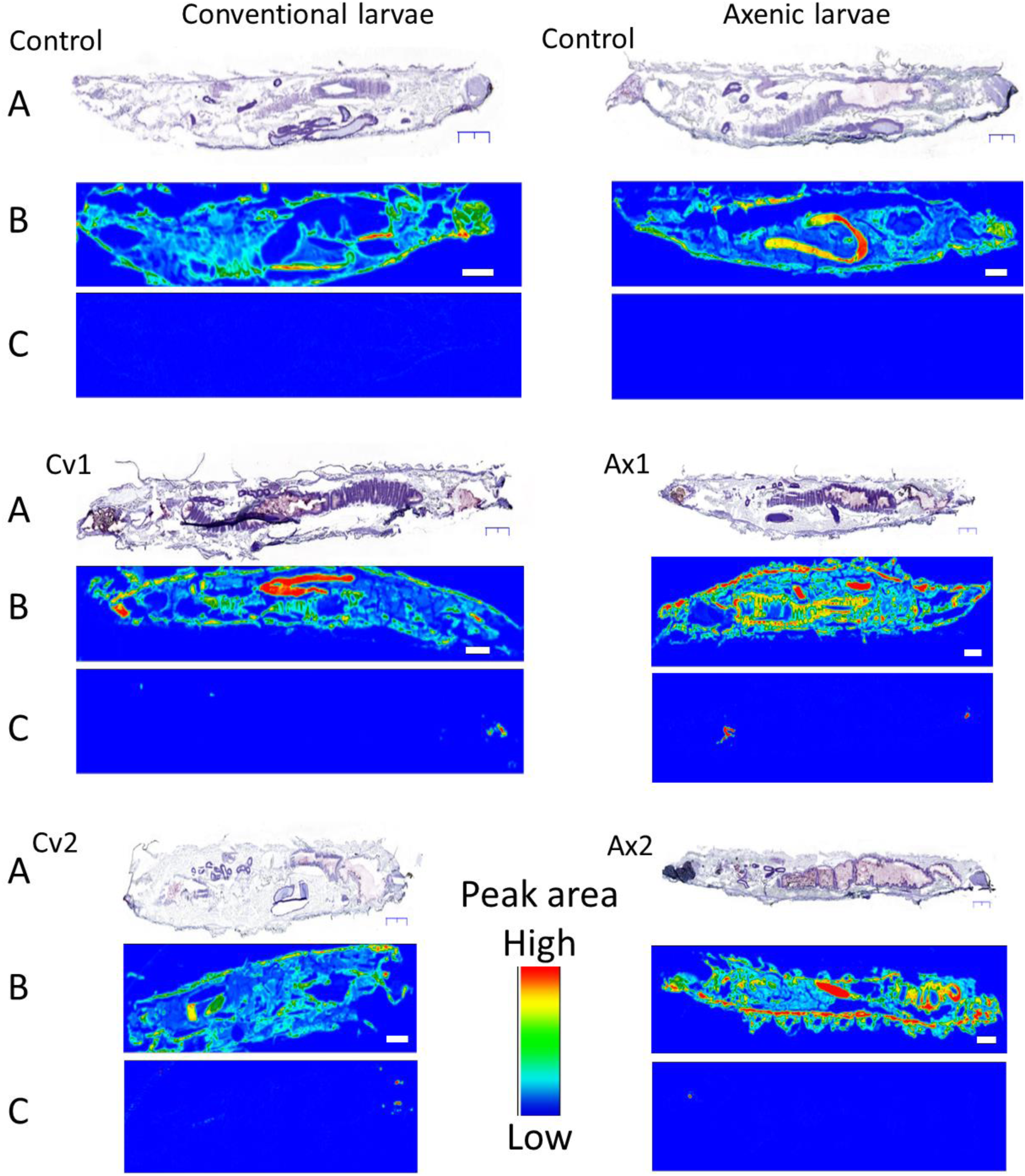
Presence of C-D signal in Gm larvae fed with PED4. Conventional (Cv) and axenic (Ax) control larvae were fed with non-deuterated PE for 3 days, and test larvae were fed with deuterated PE for 3 days (Cv1 and Ax1). Cv2 and Ax2 larvae were fed with alternating diets (see methods) over a period of 19 days. Sections were stained with cresyl-violet and observed in bright field microscopy (A) or kept unstained and analyzed by µFTIR (B and C). The IR hyperspectral images show the distribution of B) proteins (1650 cm^-1^ amide I peak area) and C) deuterated PE (2197 cm^-1^ C-D peak area). No deuterated molecules were found in the tissue of the conventional and axenic control larvae, and in the Cv1, Ax1, Cv2 and Ax2 larvae but few deuterated PE particles were detected in the mouth, gut, and rectum of both Cv and Ax larvae Spectra from such particles are shown in Figure S1F. Scale bar: 1 mm and magnification was 4X in both IR and visible micrographs.

No C-D peak was detected in the tissues of theses larvae (Cv2 and Ax2 in Figure 3).

However, particles with a C-D signal were detected in cavities of the digestive tract such as the mouth, the gut and the rectum in 6 out of 10 of the conventional and axenic larvae fed with PED4. This corresponded to the presence of micron sized PED4 particles (25-50 µm) and aggregates (up to 1000 µm). The larger PED4 particles were observed in the oral cavity and rectum. No differences were observed between axenic and conventional larvae. The particles appeared more numerous in the mouth and rectum and fewer particles were found in the gut. Since no embedding was used it is possible that some of the gut particles were lost during sectioning. While the largest particles could be detected by visual observation in the micrograph images, the C-D IR signature allowed detecting smaller particles. The spectral signature allowed confirming the chemical nature of the particles. To investigate whether smaller particles present in the gut could escape detection with hyperspectral imaging we also recorded SR-µFTIR maps and were able to detect micron-sized PE particles in the gut of the larvae (Figure S1F). We also tried to evaluate whether the PED4 found in the digestive tract was oxidised by analysing the CD_2_/CD_3_ ratio of the gut particles using the SR-µFTIR data. However, most particles were either too thick or too scattering to yield good quality spectra that could be used for such analysis.

### Biodegradation of PE by Gm larvae

The absence of PE bio-assimilation could be due to an inability of our Gm larva population to bio-degrade PE. We investigated this ability by using two methods:; detection of PED4 oxidation by µFTIR imaging of HDPE films in contact with the dissected guts of Gm larvae and analysis of the CD_2_/CD_3_ ratio of PED4 particles in the Gm larval frass by ATR-FTIR The results are detailed in the Supporting InformationWeak PE oxidation was detected in the PE films in contact with the gut of conventional Gm larvae (Figure S2) by µFTIR imaging but not with guts from the axenic larvae. Meanwhile the results did not allow to conclude on the implication of the Gm microbiota since variation were found among the replicates.

> The CD_2_/CD_3_ ratio was 15% lower for the PED4 particles 2.91±0.32) measured in the larval frass compared to PED4 films (3.42±0.40) suggesting shorter aliphatic chains in the digested PED4, thus indicating chemical modification and bio-degradation of PED4 (Figure S3).

## DISCUSSION

The capability of Gm larvae to digest and bio-assimilate PE is controversial^11,12,14,16,19,24^ and necessitates further investigation. The role of the Gm microbiota is also disputed.^12,14,18^ We therefore examined whether the Gm larvae with and without microbiota were only chewing PE or were truly able to digest and metabolize it.

Feeding experiments showed that pollen+beeswax was the optimal diet in agreement with literature.^14,17^ Early larval stages L2-L3 fed with PE lost weight, and died in 3 days (50%) to 10 days (100%). This trend is similar to results from Lou (50% death at 15 days).^18^ Kong reported 100% survival but 20-30% weight loss in PE-fed Gm larvae in 14 days.^14^ Billen reported that a LDPE diet was not sufficient for sustaining Gm larva growth.^24^ Lemoine reported a 50% weight low on a PE diet.^25^ Our results suggested that a pollen-only diet is sufficient for the survival of the larvae and that an additional source of carbon such as beeswax allows larvae to gain weight, in agreement with the literature^17,14^. L2-L3 larvae fed with pollen and PE survived 16 days and gained some weight but far less than the larvae fed with the optimal diet.

This trend was confirmed with the last stage (L6) larvae fed only with PE as they lost weight compared to larvae fed with pollen or beeswax and consumed 80 times less food than larvae with beeswax. We did not observe significant differences in feeding behaviour or weight gain between conventional and axenic larvae suggesting that the microbiota may not be important for their development and lifecycle under our condition, similar to Kong.^14^ Also for L6 larvae, the final life cycle was similar for Gm larvae fed with PE or beeswax: after 8 days the larvae pupated, the adult moths appeared 1 week later and egg production was similar. These experiments showed that PE does not have nutritional value for the Gm larvae. Larvae at early L6 stage have accumulated enough reserves to continue their lifecycle even if their diet is changed to PE alone whereas larvae at an early stage cannot survive with PE alone.

We then evaluated whether PE could be bio-assimilated by Gm larva. We used a new approach based on the µFTIR hyperspectral imaging of carbon-deuterium bonds in cryo-sections of larvae fed with deuterated PE. This method could, potentially, not only allow showing the metabolization of PE, but also help finding in which tissue PE metabolites accumulate, and what kind of biomolecules might be synthesized using PE. We first confirmed that it was possible to detect the metabolization of deuterated food in the tissues of larvae fed with perdeuterated oil mixed with pollen: C-D signal was indeed detected in most tissues and predominantly in fat body tissues after a 3-days diet in both axenic and conventional larvae. The CD_2_/CD_3_ peak area ratio was modified compared to the original oil signal showing that the oil was integrated in shorter and in longer aliphatic chains. This demonstrated that µFTIR could detect C-D signal in tissue sections from larvae that had ingested and metabolized 1-4 milligrams of deuterated food.

On the contrary, we could not detect any C-D signal in L6 larvae fed with PED4 during 3 days or during 19-21 days, although they had ingested PED4. The absence of C-D signal proved that the PE was not metabolized and integrated in the larvae bodyin substantial quantities. This should not be due to a lack of sensitivity of the spectroscopic method since the larvae ingested 2.1 mg of PED4 per larva after 3 days and 3.6 mg of PED4 per larva after 19 days (up to 5 mg) and a C-D signal was detected when larvae ingested less than 4 mg of deuterated oil. Meanwhile, PED4 micro-particles were detected in the mouth, gut and rectum of the larvae confirming that Gm larvae were able to break and masticate the PE films in smaller, micron sized, particles. Instead of metabolizing the plastic, the larvae generated micron-sized particles which could be worse for the environment and more difficult to collect than larger pieces of plastic^24^.

Yang et al.^26^ observed negligible integration of ^13^C in the body of the beetle larvae of *Tenebrio molitor* Linnaeus; fed with ^13^C polystyrene (PS), and found that large fraction of the ^13^C was integrated in the CO_2_ produced by the gut bacteria. Recently, independent results by Cassone et al.^16^ and Ren et al.^12^ both reported that bacteria isolated from the Gm larva gut were able to grow on and degrade PE into smaller compounds implicating a role of the gut microbiota. Our results showed no PE bio-assimilation in either axenic or conventional larvae. Since our Gm population was able to survive on beeswax and develope on beeswax-pollen and assimilate perdeuterated oil, it seems to possess the required enzymes to efficiently metabolize long- and short-length hydrocarbons. The absence of bio-assimilation even in conventional larvae with an intact microbiota could indicate that our Gm population lacked the bacterial strains capable of degrading PE in smaller hydrocarbons that could be processed by the gut enzymes, while these bacteria were present in the Gm population referred to in the letterature. However, we found weak oxidation in PE films in contact for 24 h with the dissected gut for both axenic and conventional larvae and a shortening of aliphatic chains in PED4 particles excreted in the larval frass indicating that a PE oxidation capability exist in our Gm. Several groups reported isolating bacterial and fungal strains capable of digesting PE from different Gm populations: Cassone reported isolating an Acinetobacter sp from Gm^16^, while Ren reported isolating an Enterobacter sp^12^, and Zhang reported isolating an Aspergillus flavus strain.^27^ Yang isolated strains of Bacillus sp and Enterobacter sp able to digest PE from the Gm related P. interpunctella larvae.^28^ The microbiota of our Gm population was analyzed by 16S rDNA sequencing and was shown to be composed of Bacteroidetes, Firmicutes, Cyanobacteria and Proteobacteria, and a few fungal species (supporting information). The Proteobacteria were mostly Enterobacteriaceae. Although Acinetobacter sp was absent, the Enterobacteriaceae strains in our Gm population suggested that its microbiota was not fundamentally different from those from other groups.^18^ Yet, Cassone suggested that bacterial communities rather than single strains would be responsible for efficient PE biodegradation. We will further evaluate the PE-degrading capacity of the Gm microbiota in our population FTIR microspectroscopy could be used to measure the integration of C-D in microcolonies of bacteria grown on PED4 films and will be tested in our follow-up work.

## ACKNOWLEDGMENTS

The authors acknowledge the Synchrotron SOLEIL for provision of synchrotron radiation facilities and FTIR microspectroscopy equipment. The authors also acknowledge the INRAE-Jouy-en-Josas histology platform (Abridge) for the cryo-sectionning and microscopy facilities and the INRAE-MICA department for the general support to CNLR and AR.

## ABBREVIATIONS

Gm: *Galleria mellonella*
PE: Polyethylene
LDPE: Low Density PE
PED4: perdeuterated polyethylene
IR: infrared
FTIR: Fourier Transform Infrared
µFTIR: FTIR microspectroscopy

## AUTHOR CONTRIBUTIONS

Agnès Réjasse (AR) initiated the study, designed the research, reared the insects, performed insect feeding experiments and participated in the writing of the manuscript.

Jehan Waeytens (JW) performed infrared microspectroscopy measurements, prepared figures and participated in the writing of the manuscript.

Ariane Deniset-Besseau (ADB) participated in the study design, and corrected the manuscript. Nicolas Crapart (NC) performed insect cryo-section and tissue coloration.

Christina Nielsen-Leroux (CNLR) participated in the study design, and corrected the manuscript.

Christophe Sandt (CS) designed the infrared microspectroscopy study, performed the infrared microspectroscopy measurements, prepared figures and wrote the manuscript.

## DECLARATION OF INTERESTS

The authors do not have any conflict of interest

## Notes

### Competing Interest Statement

The authors have declared no competing interest.

