## Supplementary tables 1-3, figures S1-S4 for "Plastic biodegradation: do *Galleria mellonella* larvae - bio-assimilate polyethylene? A spectral histology approach using isotopic labelling and infrared microspectroscopy"

### SUPPORTING INFORMATION

**Supplementary Table 1. Average food consumptions at L3 stage.**

| | **Food ingested for 16 days (mg per larva) (standard deviation)** | | | **Development after 16 days ~~at day 16~~ (except*) from L2-L3 (standard deviation)** | |
|---|---|---|---|---|---|
| **Diet** | **Pollen** | **Beeswax** | **LDPE** | **Weight gain/loss mg** | **Note** |
| **Control** | 0 | 0 | 0 | -4.5* (0.98) | All dead at day 3 |
| **Pollen** | 135.4 (11.84) | 0 | 0 | + 42.4 (11.19) | |
| **Beeswax** | 0 | 61.62 (28.17) | 0 | + 5.9 (15.4) | |
| **Beeswax + Pollen** | 187.75 (21.19) | 312.93 (28.92) | 0 | + 181.75 (38.67) | Strongest evolution |
| **LDPE** | 0 | 0 | 0.6 (0.3) | - 13.24* (8.26) | Death between day 3 and day 6 |
| **LDPE + pollen** | 158.83 (58.50) | 0 | 2.53 (2.17) | + 43.18 (30.07) | |

The food consumption is given as the average consumed weight per individual larva.

**Supplementary Table 2. Average food consumptions at L6 stage.**

| | **Food ingestion for 7 days (mg per larva) (standard deviation)** | | | **Development ~~at J7~~after 7 days from L6 (standard deviation)** | |
|---|---|---|---|---|---|
| **Diet** | **Pollen** | **Beeswax** | **LDPE** | **Weight gain (+) /loss (-) mg** | **Note** |
| **Nothing** | 0 | 0 | 0 | - 48.25 (13.15) | Weight loss |
| **Pollen** | 255.67 (21.99) | 0 | 0 | + 0.33 (7.63) | No weight change |
| **Beeswax** | 0 | 211 (67) | 0 | -5 (25.68) | No weight change |
| **Beeswax + Pollen** | 251 (32.83) | 202 (63) | 0 | + 113.25 (79.32) | Optimal diet |
| **LDPE** | 0 | 0 | 2.61 (1.69) | - 48.33 (16.43) | Weight loss |
| **LDPE + Pollen** | 179 (47.27) | 0 | 3.34 (1.24) | -6.5 (19.64) | No weight change |

The food consumption is given as the average consumed weight per individual larva.

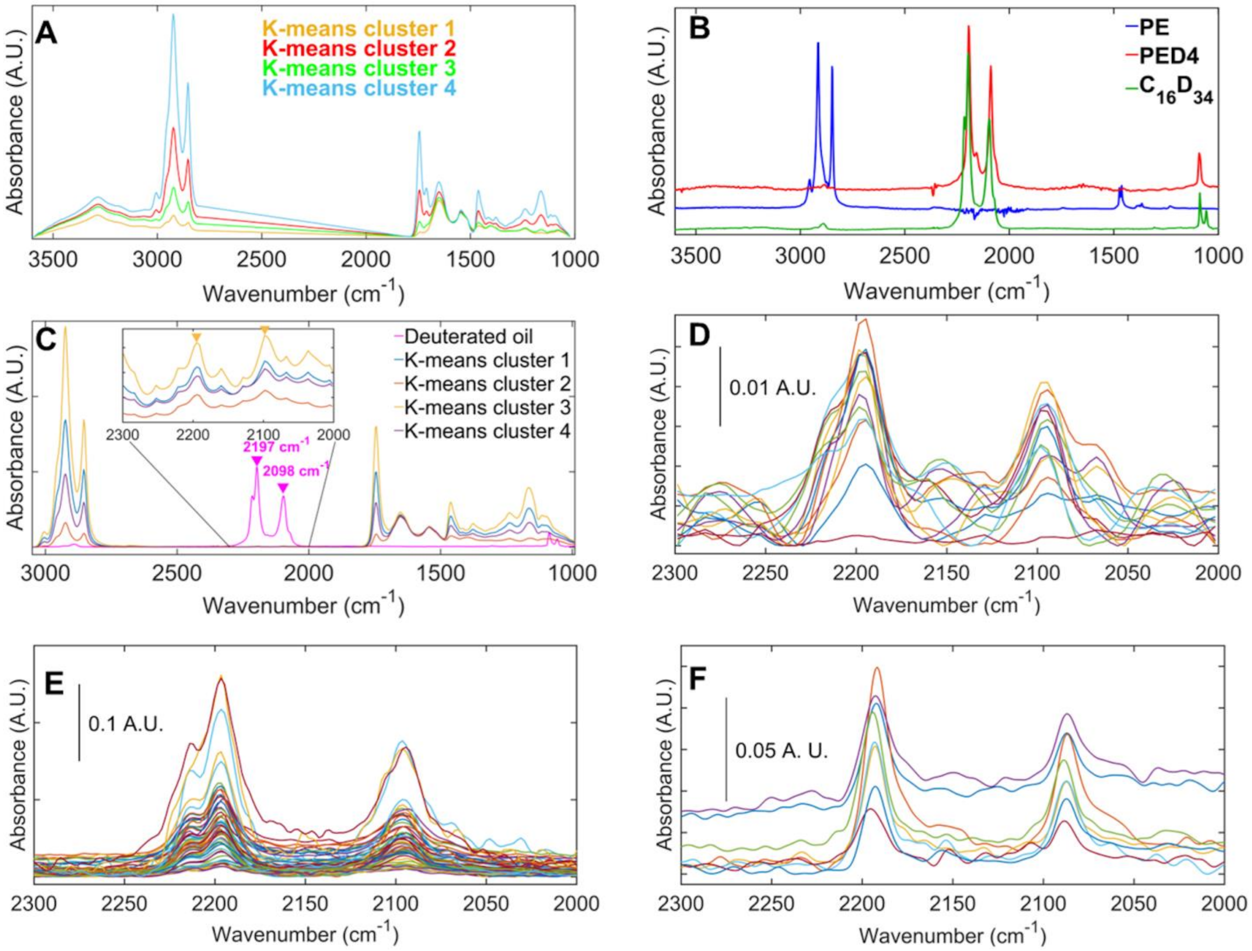


**Figure S1. Infrared spectra from larval tissues and chemicals used in this study.** A) Representative average spectra of different tissues from a control larva grouped in 4 clusters by k-means clustering. B) Infrared spectra of normal PE $(CH_2)_n$ in blue, of $C_{16}D_{34}$ perdeuterated oil in green, and of perdeuterated PE $(CD_2)_n$ in red showing the C-H peaks at circa 2920 and 2850 cm$^{-1}$ in PE, and the C-D peaks at circa 2200 and 20100 cm$^{-1}$ in the deuterated PE and the oil. Spectra were normalized and offset for clarity. C) Representative spectra from a larva fed with $C_{16}D_{34}$ oil and pollen, and $C_{16}D_{34}$ oil spectrum. The tissue spectra were grouped in 4 clusters by k-means clustering. Insert: x70 zoom on the 2300-2000 cm$^{-1}$ range containing the C-D peaks. The fatter tissues contain more C-D. D) IR spectra extracted from the C-D rich region in Figure 2 along the line shown in Figure 2K. Only the 2250-2050 cm$^{-1}$ C-D peak region of the IR spectra is shown. E) IR spectra from C-D rich regions of a Gm larva fed 72h with $C_{16}D_{34}$ oil measured with a confocal microscope coupled to the synchrotron radiation source; the C-D peaks at 2197 and 2098 $cm^{-1}$ reached absorbance values one order of magnitude higher than those measured with the hyperspectral imaging microscope, increasing measurement sensitivity and accuracy. F) Synchrotron-μFTIR spectra from PE microparticles in the gut of a PED4 fed larva.

***Bio-degradation of PE films by dissected gut of Gm larvae***

Since we could not detect any PE oxidation in the gut of the larvae, we then asked the question whether PE could be directly oxidised by the gut of the Gm larvae as reported by others[11]. This approach was criticized in Weber et al. (2017)[17] due to possible confusing factors such as contaminations and misinterpretations. Here we decided to use µFTIR hyperspectral imaging to study whether Gm gut content with or without microbiota could oxidize PE films. This approach allowed finely differentiating between regions where the PE film was oxidized and regions were the PE film was contaminated by residues of the larval gut. Guts from conventional and axenic Gm larvae were dissected and deposited on the surface of small pieces of an 8 µm thick HDPE (High density Polyethylene) plastic bag and incubated at 27°C in humid atmosphere for either 24 hours or 21 days (Figure S2A). The HDPE plastic bag pieces were then cleaned following a specifically developed protocol to eliminate gut residues (see the Experimental section at the end).

IR hyperspectral images of the HDPE were recorded in transmission mode. PE was characterized by its methylene peak at 1465 $cm^{-1}$ and PE oxidation by the carbonyl peak around 1735 $cm^{-1}$. The carbonyl/methylene peak area ratio was used to image the distribution of PE oxidation in the PE films in contact with the Gm larva gut. The presence of eventual gut residues was detected by measuring the presence of amide I (1650 $cm^{-1}$) and amide II (1545 $cm^{-1}$) bands from proteins. Clean pieces of HDPE bags (no contact with Gm guts) were used as control. Figure S2B shows representative spectra from the different plastic bags. The orange spectrum came from a control bag that was found to be naturally oxidised and showed a strong carbonyl peak with several shoulders evidencing multiple oxidation levels. It is noteworthy that the level of oxidation in that control bag was higher than in any of the treated bags by several orders of magnitude.

Even in the control bags we could detect weak carbonyl peaks at some discrete positions by using the hyperspectral images built from the carbonyl/methylene ratio (Figure S2C). This could arise from natural PE oxidation but also from the presence of other molecules such as polymer additives (plasticizers, anti-oxidants…).

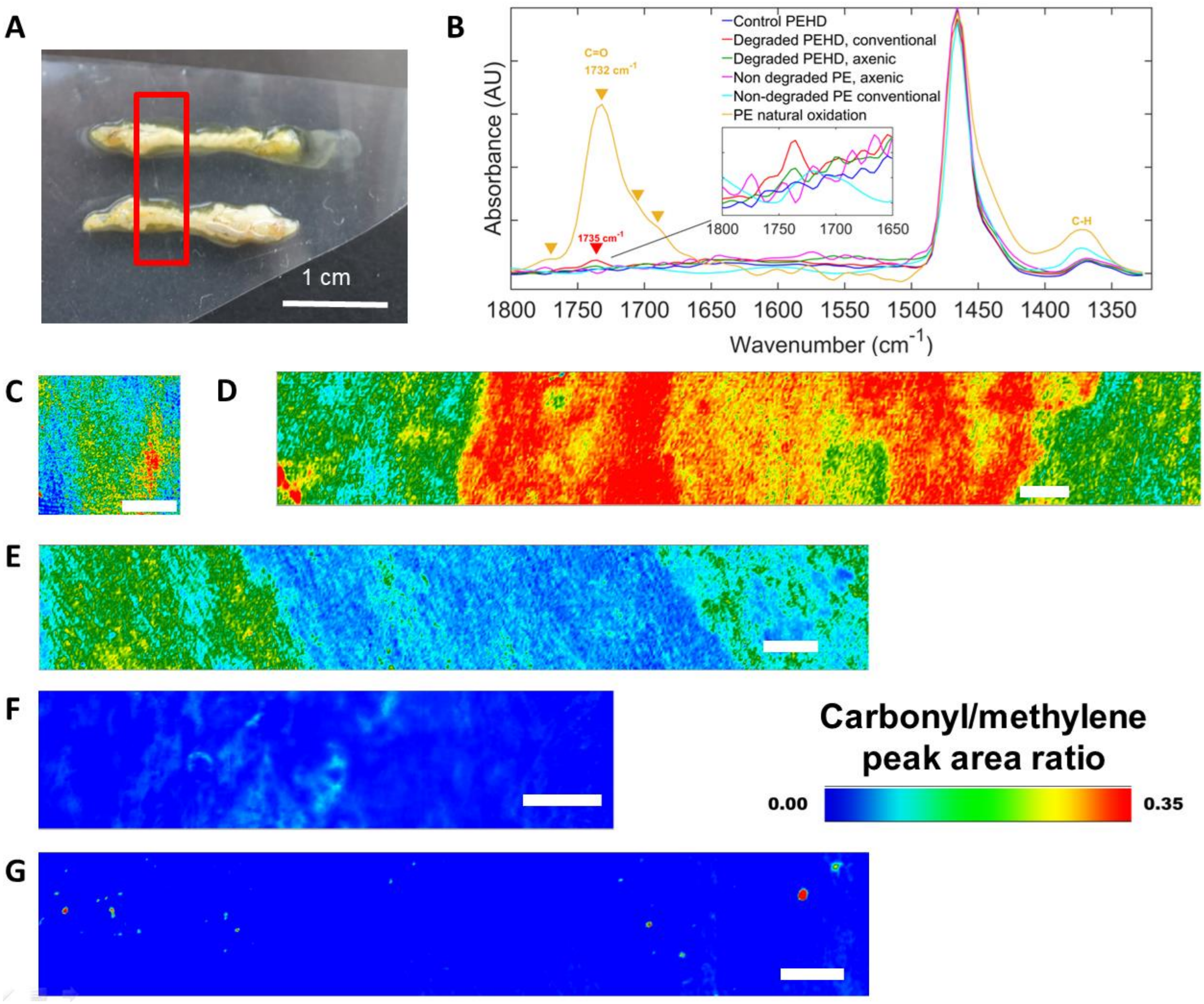


**Figure S2. µFTIR imaging analysis of the degradation of HDPE films in contact with Gm larva gut.** PE films were incubated at 27°C in humid atmosphere with the dissected gut of conventional or axenic larva deposited at the surface of the film, and spectral images of the PE films were recorded after 24 h or 21 days of contact. Hyperspectral images were generated by plotting the carbonyl/methylene (C=O/C-H) peak area ratio using the C=O peak at 1735 $cm^{-1}$ and the C-H peak at 1370 $cm^{-1}$ after water vapour subtraction (when necessary). Micrograph scale bar: 1 mm. A) Optical view of the dissected larva guts deposited on pieces of HDPE bags. B) Spectra of PE films in the 1300-1800 $cm^{-1}$ range. Non-oxidized film (blue), ambient-air oxidized film (orange). The air oxidized film presents several peaks characteristic of various C=O moieties between 1650 and 1780 $cm^{-1}$ and is dominated by the C=O from esters at 1732 $cm^{-1}$. Representative carbonyl/methylene spectral images of the PE films are shown in images C to F. A weak C=O peak at 1735 $cm^{-1}$ characteristic of oxidation is observed in some but not all of the films. C) control PE film showing regions with various oxidation levels. D) PE film degraded by the gut of a conventional Gm larva for 24h. E) PE film degraded by the gut of an axenic Gm larva for 24h. F) PE film degraded by the gut of a conventional Gm larva for 21 days. G) Image of the amide I band on a PE film degraded by the gut of an axenic Gm larva for 24h showing traces of protein and fat (green and red spots). The intensity scale of image G is 0.5-3.0 absorbance unit.

A weak C=O peak at 1735 $cm^{-1}$ was detected in about two-third of the films exposed to the gut extract (Figure S2D-F) for both conventional and axenic samples but was undetectable in the other films. In the carbonyl/methylene peak area ratio images, the location of the carbonyl peak matches with the location of the gut on the film (Figure S2D, S2E). The carbonyl/methylene peak area ratio is reported in Supplementary Table 3). It was several times higher in the oxidized part of the plastic than in the non-oxidised regions or in the control film. However, the C=O peak was not detected or was lower than the control in some of the treated samples (conventional and axenic); and could barely be detected in the two samples incubated during 21 days (Figure S2F and Supplementary Table 3).

**Supplementary Table 3.** Oxidation level of PE by the gut of Gm larvae (Cv, conventional and Ax, axenic).

| Larva # | Treatment | Duration (day) | HDPE oxidation CO:CH ratio | Change relative to control | Notes |
|---|---|---|---|---|---|
| **T1** | Control | - | 0.54 | - | Discrete oxidation |
| **Cv I1** | Gut + microbiota | 3 | 0.64 | 1.18 | Strongest |
| **Cv I2** | Gut + microbiota | 3 | 3.10 | 5.70 | Contamination with NaH(CO3) |
| **Cv I3** | Gut + microbiota | 3 | 0.30 | 0.55 | Discrete oxidized spots |
| **Cv I4** | Gut + microbiota | 21 | ? | - | Strong water vapour |
| **Cv I5** | Gut + microbiota | 21 | 0.03 | 0.04 | Interference fringes |
| **Ax I1** | Gut | 3 | 0.05 | 0.09 | Contamination with unknown carbohydrate |

| | | | | | |
|---|---|---|---|---|---|
| **Ax I2** | Gut | 3 | ? | - | Contamination with NaH(CO3) |
| **Ax I3** | Gut | 3 | 0 | - | No detectable oxidation |

With the use of a rigorous cleaning protocol, over 90% of the film was clean of fats and proteins but we could detect the presence of gut residues at discrete positions of some PE films (FigureS2G), and also other contaminations originally present in the bags such as carbohydrates and $NaH(CO_3)$.

This experiment revealed several confusing factors such as water vapour signal, interference fringes, and residues from gut proteins and fats that obscured the detection of PE oxidation since the carbonyl peak was weak in all samples. The O-H bending mode from water vapour also strongly absorbs at 1735 $cm^{-1}$. Although the IR microscope was continuously purged with nitrogen, the water vapour signal was often found to be of the same amplitude as the C=O peak in the oxidized HDPE and a water vapour subtraction step had to be performed to confirm whether any oxidation was detected. Furthermore, even after thorough cleaning procedure was applied to HDPE bags to remove the gut tissue, discrete spots with residual proteins and lipids were detected in some samples. The C=O stretching mode from esterified lipids absorb at 1740 $cm^{-1}$. PE samples that are not correctly cleaned will be contaminated by fats whose stronger carbonyl band will totally obscure the weak carbonyl band from oxidized PE. These confusing factors limited both the sensitivity and specificity of the protocol. Although our findings suggested that PE might be weakly oxidised by the extracts of Gm larva gut and that the carbonyl/methylene ratio was stronger in the conventional than in the axenic cases, oxidation signals were weak and barely above detection threshold. Following this protocol, w~~W~~e thus cannot firmly conclude about the implication of the gut microbiota in the oxidation of the PE.

***Material and methods***

Gm gut dissection protocol: 6 conventional and 6 axenically reared larvae were dissected, the larvae were cut longitudinally from the ventral side, and the skin spread with pins, the fat body tissues were removed, and the digestive tube was removed from the mouth to the anus. The digestive tubes were opened longitudinally so that the inside of the gut could be placed in contact with the PE films.

HDPE plastic bags were cut in 3 cm by 7 cm triangular pieces. The HDPE pieces were sterilized on each side by UV light at 254 nm for 10 min. Two complete digestive tubes were placed on each of the HDPE pieces, then deposited in 6 Petri dishes and incubated at 27°C in humid atmosphere for either 24 hours or 21 days.

After the incubation of the digestive tubes on the HDPE, the pieces were cleaned by a specifically developed protocol: rinsed and shaken in 2 ml of water, followed by another bath in 5 ml of water at 60 ° C, then rinsed in 50% ethanol at 60°C to remove fat, then washed 5 times for 5 minutes in NaOH 1M at room temperature in an ultrasonic bath, and finally rinsed in distilled water and dried at room temperature.

***Change in the $CD_2/CD_3$ ratio in PED4 particles measured in larval frass by ATR-FTIR***

The infrared spectra of the larval frass fed with PE or PED4 were measured by the ATR-FTIR technique. The presence of PED4 particles in the frass was evidenced by the C-D peaks between 2100 and 2200 $cm^{-1}$. Larva fed normal PE did not present any C-D peaks. Since the PED4 signal was low (0.02-0.008 A.U. at 2191 $cm^{-1}$ for the stronger $CD_2$ peak) and due to the strong overlap of the $CD_2$ and $CD_3$ peaks, we used curve-fitting to extract the peak parameters. The C-D region of the spectra was fitted by 5 Gaussian/Lorentzian curves: 2191 ($\nu_{as}$ $CD_2$), 2175 (HCD), 2156 ($\nu_{as}$ Fermi resonance), 2086 ($\nu_s$ $CD_2$), 2066 ($\nu_s$ $CD_3$) $cm^{-1}$. The $CD_2/CD_3$ peak height ratio (RCD) was used to evidence PED4 oxidation and was computed for both the symmetric ($\nu_s$) stretching vibrations. The antisymmetric $CD_3$ stretching vibrations between 2200 and 2250 $cm^{-1}$ were barely noticeable.

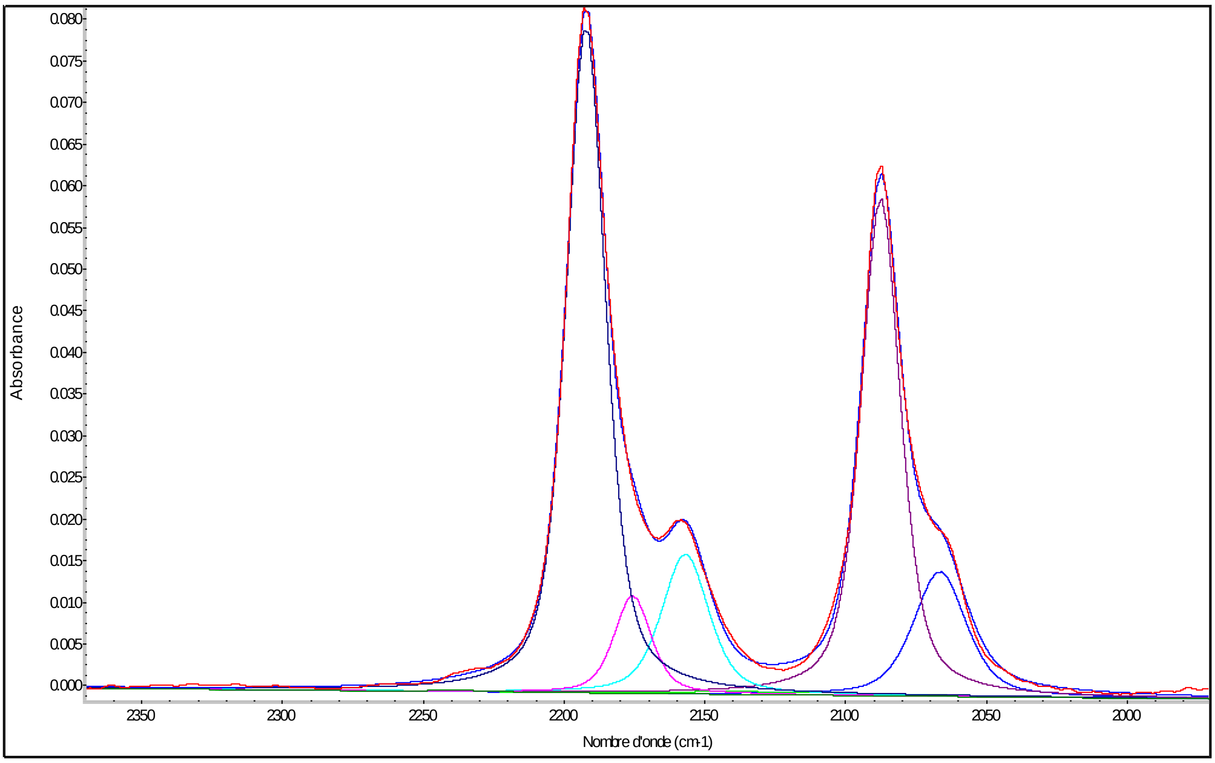


**Figure S3. Curve-fitting of the C-D region of PED4 measured in the frass of a conventional larva.** The PED4 peaks were fitted by components at 2191 ($\nu_{as}$ $CD_2$), 2175(HCD), 2156(Fermi Resonance), 2086 ($\nu_s$ $CD_2$), 2066 ($\nu_s$ $CD_3$) $cm^{-1}$.

Control PED4 films measured by the same method had a RCD of 4.46±0.29. The PED4 in frass had a lower RCD of 3.62±0.29, a 19% difference. Although the values were not found statistically different by a t-test (presumably a statistical oddity due to the low number of control PED4 films), we suggest that the change in $CD_2/CD_3$ peak height ratio indicated that the PED4 found in the frass had somewhat been processed and its aliphatic chain length had been reduced.

***Material and methods***

Larval frass were collected and stored at -80°C until measurement. Spectra were measured with a Perkin Elmer Spectrum 100 spectrometer (Perkin-Elmer, Les Ulis, France) equipped with a one millimetre diamond ATR crystal and DTGS detector. The spectra were recorded at 4 $cm^{-1}$ resolution between 400 and 4000 $cm^{-1}$ with 16-32 co-added scans.

The spectra were fitted with the Peak Resolve module of Omnic 9.2 software. The 1970-2370 $cm^{-1}$ region containing 208 points was fitted with 5 Gaussian/Lorentzian peaks for a total of 35 free parameters (position, width, intensity, Gaussian factor; baseline).

**Analysis of Gm larvae microbiota by 16S rRNA sequencing**

*Experimental*

To analyse the microbiota of *Galleria mellonella*, L6 larvae (average weight 200 mg) were sampled from the Jouy-en-Josas rearing (fed with pollen and beeswax) and left without food for 24h at 27°C. Feces were collected, larvae were dissected and the whole guts were removed and stored at -80 °C as were the feces.

DNA extraction from Galleria mellonella gut and feces: The weight of guts and feces used for DNA extraction were respectively 30-35 mg and 10-15 mg. The DNA extractions were carried out with the "Powerlyser PowerSoil DNA isolation kit" (MO BIO laboratories, inc.) according to the manufacturer's instruction with slight modifications as follows. After the addition of solution C1, samples were incubated at 65°C with stirring at 900 rpm for 10 min and high speed centrifugation for 1 min at 20,000g was performed after the homogenization step, using FastPrep 24 instrument (MP Biomedicals) at 4m/s twice for 40s with a 5 min break at 4°C between each run. The DNAs were quantified with a Nanodrop 2000 (Thermo Scientific). The DNA concentrations were between 30 and 40 ng/µL per gut and the ratios 260:280 nm and 260:230 nm were >1.8. For the feces the ratios were >1.8 and the concentration was between 15 and 20 ng/µL.

Amplicon libraries, Illumina Miseq sequencing and analysis:

Amplicon libraries were constructed following two rounds of PCR amplification. The first amplification of the~450-bpV3- V4 hypervariable regions of the bacterial 16S rRNA gene was performed with the primers V3F (5'-ACGGRAGGCWGCAG- 3') and V4R (5'-TACCAGGGTATCTAATCCT-3') as described in Poirier et al.[21] Amplicon size, quality and quantity were checked on a DNA 1000 chip (Agilent Technologies, Paris, France). The purification and quantification of the second Illumina MiSeq PCR was made by Genotoul (Toulouse-France) as well as the Illumina Miseq sequencing. Quality filtering, definition of OTUs and taxonomic assignment was made by Stephane Chaillou (INRAE-MICALIS, FME team) using FROGS pipeline (Find Rapidly OTU with Galaxy Solution).

*Results*:

The bacterial microbiota of the G. *mellonella* population analyzed by 16S rRNA sequencing, showed a composition largely dominated by two Enterococcus species, E. *mundtti* and E. *gallinarum*.  Indeed the two species represent 98 % of the reads, in both feces and whole dissected guts (figure S4). In the remaining 2 % of the reads a few other species were found: other gram positive firmicutes as *Enterococcus faecalis* and *Lactobacillus kunkeei*, and the gram-negative *Enterobacter arsenophonus*. This indicates that the here studied *Galleria mellonella colony* is not very riche in bacterial diversity, and it is similar to the study of Lou et & al. (2020).

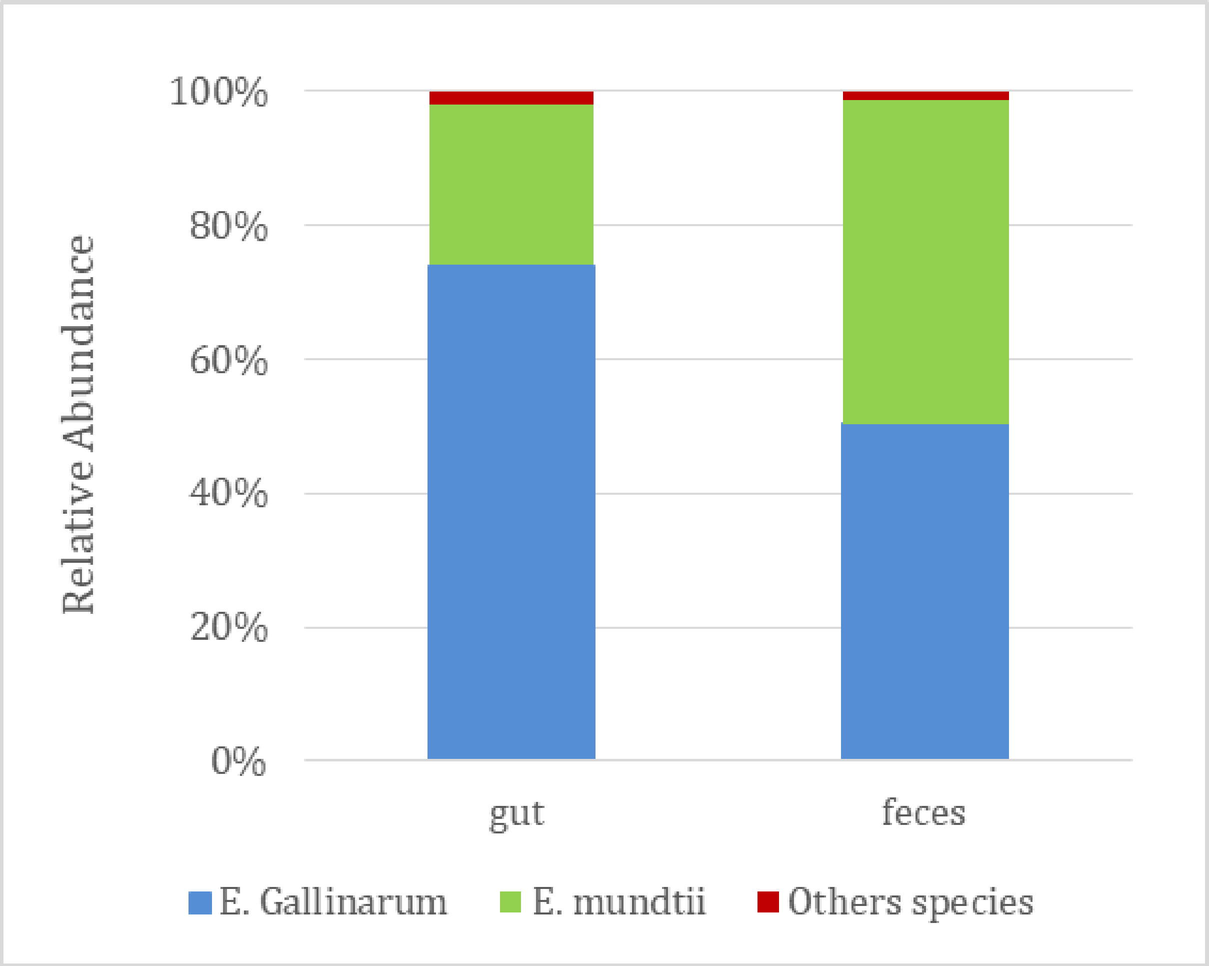


**Figure S4: Relative abundance of bacterial species.** Abundances following reads from MiSeq sequencing of 16S rDNA obtained from dissected guts (three replicates) and feces (six replicates) *from Galleria mellonella* larvae. 100% is the total number of reads.
